# Focal acetylcholinergic modulation of the human midcingulo-insular network during attention: Meta-analytic neuroimaging and behavioral evidence

**DOI:** 10.1101/2023.09.20.558618

**Authors:** Sudesna Chakraborty, Sun Kyun Lee, Sarah M. Arnold, Roy A.M. Haast, Ali R. Khan, Taylor W. Schmitz

## Abstract

The basal forebrain cholinergic neurons provide acetylcholine to the cortex via large projections. Recent molecular imaging work in humans indicates that the cortical cholinergic innervation is not uniformly distributed, but rather may disproportionately innervate cortical areas relevant to supervisory attention. In this study, we therefore reexamined the spatial relationship between acetylcholinergic modulation and attention in the human cortex using meta-analytic strategies targeting both pharmacological and nonpharmacological neuroimaging studies. We found that pharmaco-modulation of acetylcholine evoked both increased activity in the anterior cingulate and decreased activity in the opercular and insular cortex. In large independent meta-analyses of non-pharmacological neuroimaging research, we demonstrate that during attentional engagement these cortical areas exhibit (1) task-related co-activation with the basal forebrain, (2) task-related co-activation with one another, and (3) spatial overlap with dense cholinergic innervations originating from the BF, as estimated by multimodal PET and MR imaging. Finally, we provide meta-analytic evidence that pharmaco-modulation of acetylcholine also induces a speeding of responses to targets with no apparent tradeoff in accuracy. In sum, we demonstrate in humans that acetylcholinergic modulation of midcingulo-insular hubs of the ventral attention/salience network via basal forebrain afferents may coordinate selection of task relevant information, thereby facilitating cognition and behavior.

## Introduction

Acetylcholine (ACh) is a neurotransmitter that plays a critical role in modulating neural activity and information processing in the brain. The primary source of neocortical, hippocampal and amygdalar ACh originates from the cholinergic neurons in the basal forebrain (BF), which consists of several subcortical nuclei situated adjacent to the hypothalamus (Mesulam and Geula 1988; Zaborszky *et al*. 2015; Woolf 1991). Individual BF cholinergic neurons have immense axonal projections, which can branch >1000 times before reaching their synaptic targets in the cerebrum (Wu *et al*. 2014).

Most of what we understand about the influence of ACh on cortical function comes from research in non-human animal models. Optogenetic and electrophysiological studies demonstrate that cholinergic BF neurons emit precisely timed ACh signals in response to the detection of novel or salient sensory stimuli, as well as task-relevant outcomes such as rewards and errors, at the millisecond timescale (Hangya *et al*. 2015; Pinto *et al*. 2013; Guo *et al*. 2019; Laszlovszky *et al*. 2020; Harrison *et al*. 2016; Tu *et al*. 2022; Bennett *et al*. 2012; Letzkus *et al*. 2011). Advances in biosensor strategies indicate that ACh signaling can exhibit spatial specificities ranging from the macroscopic scale of different brain regions (Teles-Grilo Ruivo *et al*. 2017) down to the micron scale of cortical ensembles and receptive fields (Jing *et al*. 2020; Sethuramanujam *et al*. 2021). These discoveries have revealed ‘wired’ transmission modes of ACh signaling, which in turn have necessitated revisions to our understanding of its role in cortical function (Sarter and Lustig 2020; Sarter *et al*. 2009; Schmitz and Duncan 2018; Záborszky *et al*. 2018). At the network level, wired ACh neurotransmission may hierarchically integrate cortical areas to a “read-in” mode, while suppressing cortical consolidation, or “read-out”, of memory (Hasselmo and McGaughy 2004). High levels of ACh release thus appear to set local and global neural dynamics that are optimal for attention, encoding and learning.

We recently demonstrated with multimodal MRI and PET data acquired from humans that cortical regions including the dorsal anterior cingulate, anterior insula and frontal operculum express molecular, structural and functional markers of BF cholinergic innervation which are disproportionately higher than other cortical regions (Chakraborty *et al*. 2023) (Figure 1A). Further analysis revealed that these regions closely overlap with hubs of the ventral attention network (Yeo *et al*. 2011; Alves *et al*. 2022; Corbetta *et al*. 2008; Vossel *et al*. 2014) and salience network (Seeley *et al*. 2007; Seeley 2019). The anatomical core of the salience and ventral attention networks comprises a set of midcingulo-insular cortical hubs, according to recent taxonomies of cortical networks (Uddin *et al*. 2019). The midcingulo-insular network mediates switching between the default mode and central executive networks (Sridharan *et al*. 2008; Uddin 2015), consistent with a domain-general role in coordinating attentional resources throughout the brain (Vossel *et al*. 2014; Downar *et al*. 2000; Menon and Uddin 2010). The phylogenetic cortical homologues of a midcingulo-insular network are also observed in the intrinsic cortical connectome of the mouse (Mandino *et al*. 2022; Sforazzini *et al*. 2014), suggesting evolutionary conservation of a core attention system for saliency, switching, and control (Figure 1B). In mouse and human, these systems receive dense inputs from highly branched BF cholinergic neurons (Chakraborty *et al*. 2023; Li *et al*. 2018).

**Figure 1:**
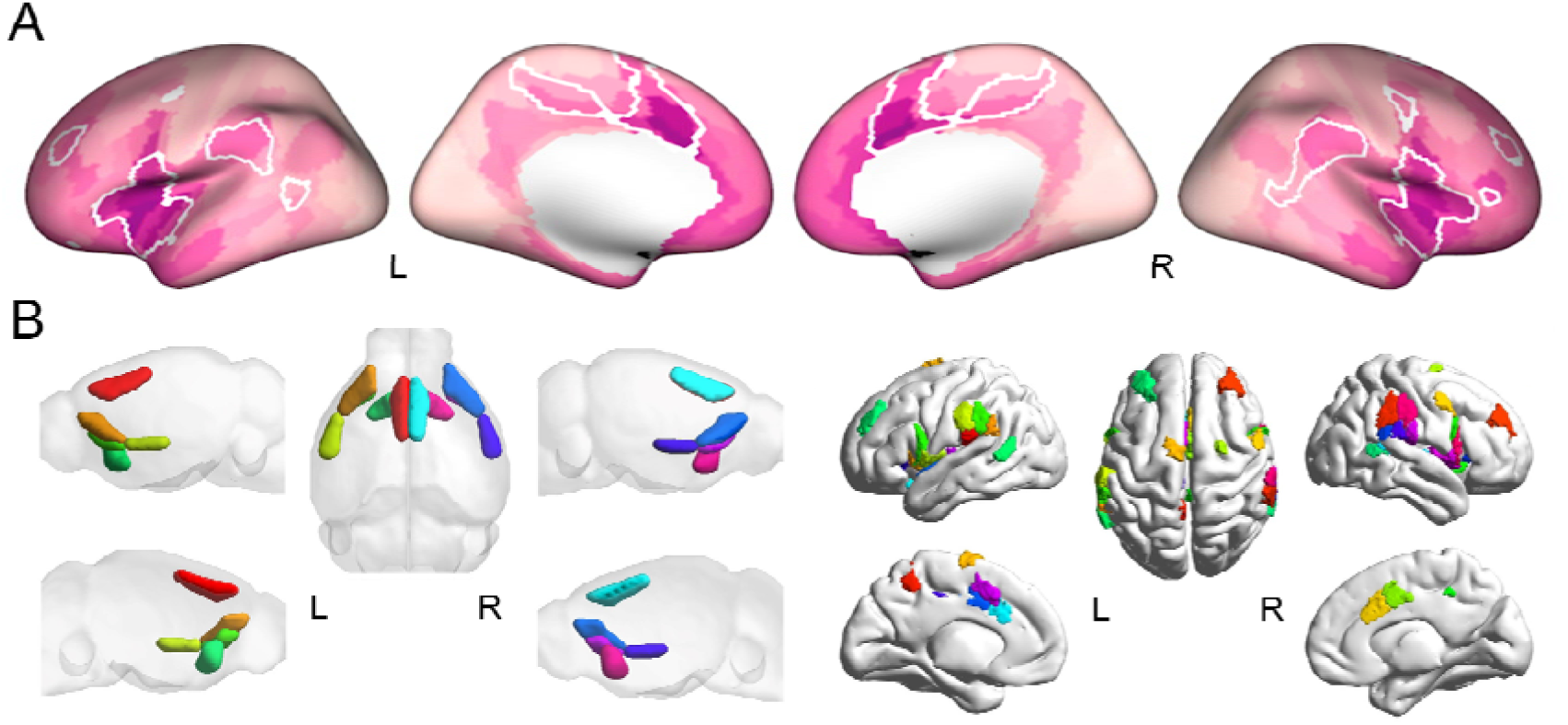
Homologous midcingulo-insular hubs of the ventral attention/salience networks in the human and mouse brain. (A) Surface rendering displaying the human ventral attention network (cortical hubs delineated by white borders) overlaid on a multimodal map of the BF connectome (Chakraborty et al. 2023), estimated from cortical VAChT concentration ([18F] FEOBV PET), BF structural (diffusion-weighted MRI) and functional connectivity (resting-state fMRI), geodesic distance from BF, and cortical myelin (T1w/T2w MRI). Areas with the darkest color exhibit higher VAChT, higher structure-function detethering in measures of BF connectivity, shorter geodesic distances to BF, and lower cortical myelin. (B) The ventral attention/salience networks exhibit homologous midcingulo-insular cortical architecture and function in mouse (left) and human (right); this figure is adapted with permission from Xu et al (2022).

However, in humans the direct link between ACh signaling and attention has proven challenging to study experimentally. Biomarkers that can reliably measure the time-varying functional activity of the cholinergic system with cell type specificity are currently feasible in non-human animals only. One strategy for overcoming this obstacle is to measure task-related brain activity with neuroimaging techniques such as fMRI or PET while participants are administered pharmacological interventions targeting cholinergic function. Drugs which alter cholinergic function fall into one of two general categories: ACh agonists and ACh antagonists. For reviews, see Bentley et al (2011) and Sutherland et al (2015). From this work, a complex and sometimes contradictory picture of the relationship of ACh with task-related activations and task performance has emerged. For instance, pharmacological activation of ACh alters task-related brain activation in areas associated with attentional orienting, including cortical hubs of the ventral attention/salience network. However, studies also report many other patterns, including increased task-related activity in parietal and sensory areas, and decreased activity in cortical midline areas. Concurrent effects of ACh activation on behavioral performance also vary from study to study. While some studies report performance facilitation under ACh activation, e.g., faster responses and higher accuracy for visuospatial target detection, others report negligible effects.

Meta-analysis is a powerful tool for improving consensus across empirical research studies. Well-curated meta-analytic inclusion criteria can provide focused integration of research examining a target population with a common set of experimental factors. Moreover, meta-analysis provides quantitative integration of observations across studies. The resulting increase in sample size leads to increased statistical power to detect smaller effects, such as a common pattern of task-evoked brain activation or behavior, that may have been missed in individual studies. The pharmacological neuroimaging literature on ACh is broad. Studies have examined populations ranging from cognitively normal to those with conditions thought to affect central cholinergic integrity, including schizophrenia, substance abuse disorders, autism, and Alzheimer’s disease. Moreover, many studies use tasks which are not designed to engage directed attention, such as passive viewing, or use task-free resting-state fMRI paradigms. Heterogeneity in these experimental factors poses a major challenge to synthesis of coherent patterns across studies.

We therefore performed several meta-analyses to examine the influence of ACh and directed attention on human brain function and behavior. Our strategy combined (1) focused meta-analysis on a well curated sample of pharmacological neuroimaging studies with (2) discovery and validation meta-analyses on larger independent samples of non-pharmacological neuroimaging studies (See Figure 2). To address heterogeneity in experimental factors, the pharmacological imaging meta-analysis focused on experiments that met the following three criteria: (1) employed ACh agonists with placebo control; (2) report explicit task comparisons of high versus low attentional demand conditions; (3) assessed cognitively normal younger adults (<50 years). Our meta-analyses of non-pharmacological imaging studies provided data-driven discovery of cortical areas that co-activate with the BF during task engagement, and validation that cortical areas modulated by ACh and attention (identified in the meta-analysis of pharmacological neuroimaging) also co-activate with one another during task engagement. Sample sizes in both the discovery and validation meta-analyses exceeded 80 unique neuroimaging studies, comprising >1300 individuals. Lastly, we performed to our knowledge the first behavioral meta-analysis of ACh pharmacological neuroimaging studies to determine if cholinergic modulation impacts response latency and accuracy measures of attentional performance.

**Figure 2:**
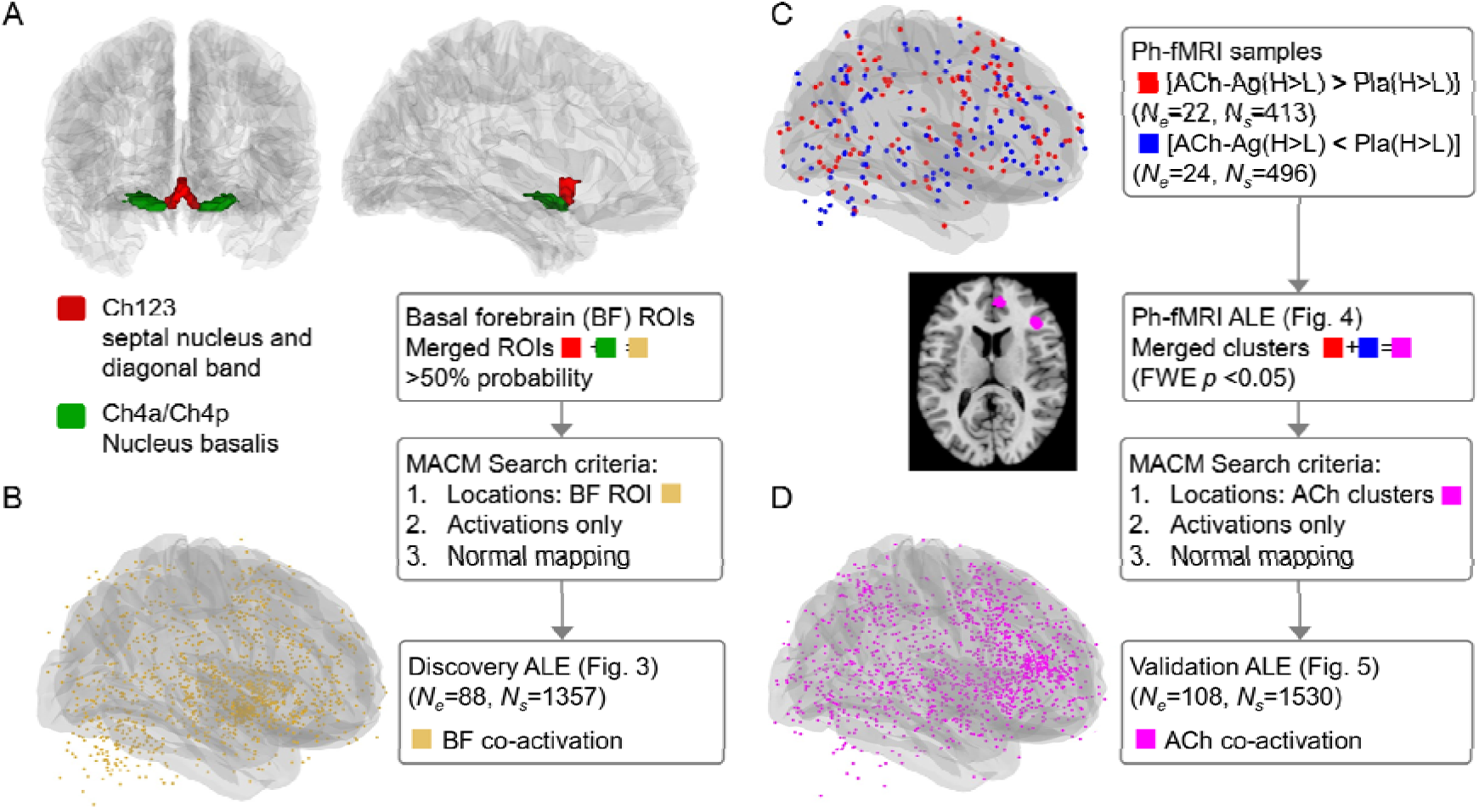
Neuroimaging meta-analysis strategies. (A) A probabilistic atlas of the Ch123,Ch4a/4p BF nuclei (Zaborszky et al. 2008). (B) The BF atlas was used as a seed region for meta-analytic connectivity mapping (MACM). Additional search filters restricted our sample to neuroimaging studies reporting task activation in cognitively normal adults. The MACM localized coordinates for brain areas that co-activate with the BF under attentional engagement (gold dots in semi-transparent render). These coordinates were submitted to a discovery activation likelihood estimation (ALE) analysis (Figure 3). (C) We identified placebo (Pla) controlled pharmacological neuroimaging studies that: (1) employed cholinergic agonists (ACh); (2) report explicit task comparisons of high versus low (H>L) attentional demand condition; (3) assessed cognitively normal younger adults (<50 years). ALE was performed separately on Drug x Task interactions where ACh either increased activation under ACh and attention (red dots: [ACh-Ag(H>L)]>[Pla(H>L)], or decreased activation under ACh and attention (blue dots: [ACh-Ag(H>L)]<[Pla(H>L)]; Figure 4. (D) Suprathreshold ALE clusters for activation increases and decreases were merged into a seed region (magenta dots in semi-transparent render) for MACM. The MACM localized coordinates for brain areas that co-activate with the ACh modulated cortical regions under attentional engagement. These coordinates were submitted to validation ALE analysis (Figure 5).

Altogether, we demonstrate that pharmacological activation of ACh alters distributed patterns of midcingulo-insular brain activity during directed attention. These regions exhibit strong co-activation with the BF and with one another during task engagement, close spatial alignment with a multimodal map quantifying the gradients of cortical cholinergic innervation originating from the BF, and are associated behavioral markers of facilitated attentional performance.

## Materials and Methods

### Literature Search and Selection Criteria for Pharmacological neuroimaging studies

We conducted a literature search on the PubMed database (https://pubmed.ncbi.nlm.nih.gov) to identify pharmacological functional imaging studies of potential relevance to the current hypothesis up to April 2023. The search terms used were adopted from Bentley et al. (2011) and contained the following combinations of keywords: [cholinergic OR acetylcholine OR nicotine OR scopolamine OR cholinesterase OR smoking OR varenicline] AND [functional imaging OR fMRI OR PET]. Meta-analytic review papers (Sutherland *et al*. 2015) and reference lists were also examined to retrieve additional relevant studies. Following the initial screening of titles and abstracts, the selected articles underwent further screening based on predefined eligibility criteria. Studies were included in the meta-analysis if they (1) used fMRI or PET (2) reported activation data obtained from healthy, neurologically intact, non-elderly (i.e. mean age < 50 years) participants; (3) involved cholinergic manipulation of participants by either pharmacological administration or cigarette smoking in a placebo-(or abstinence) controlled within-subjects or between-subjects design; (4) employed tasks with at least two levels of cognitive load/demand; (5) reported activation coordinates for brain regions that demonstrated interaction between drug and task; and (6) reported peak activation coordinates in standardized stereotaxic space (i.e. either Montreal Neurological Institute (MNI) or Talairach space). Studies investigating functional connectivity were excluded from this meta-analysis. From our meta-analytic search criteria above, we determined that the use of ACh antagonists such as atropine and scopolamine is less common in the neuroimaging literature (*N_studies_*=8, *N_subjects_*=126). The sample size for antagonist studies which met our search criteria is considerably less than 17 which has been suggested as a lower bound for well powered ALE meta-analyses (Yeung *et al*. 2023; Eickhoff *et al*. 2017). While we did not perform statistical inference on the ACh antagonist studies, ALE configuration files for their reported coordinates are available at our github (see below).

### Activation Likelihood Estimation (ALE)

Relevant coordinates from the selected studies were extracted and converted into MNI coordinates, where necessary, using the Lancaster transform included in GingerALE v.3.0.2 (Lancaster *et al*. 2007). For ACh agonists, activation foci were separated according to their pattern of task-evoked activity under ACh modulation and attention. We then ran two meta-analyses: (1) Activation increases under ACh modulation and attention [ACh-Ag(H>L)>Pla(H>L)] and (2) Activation decreases under ACh modulation and attention [ACh-Ag(H>L)<Pla(H>L)]. See Tables 1 and 2.

**Table 1.**
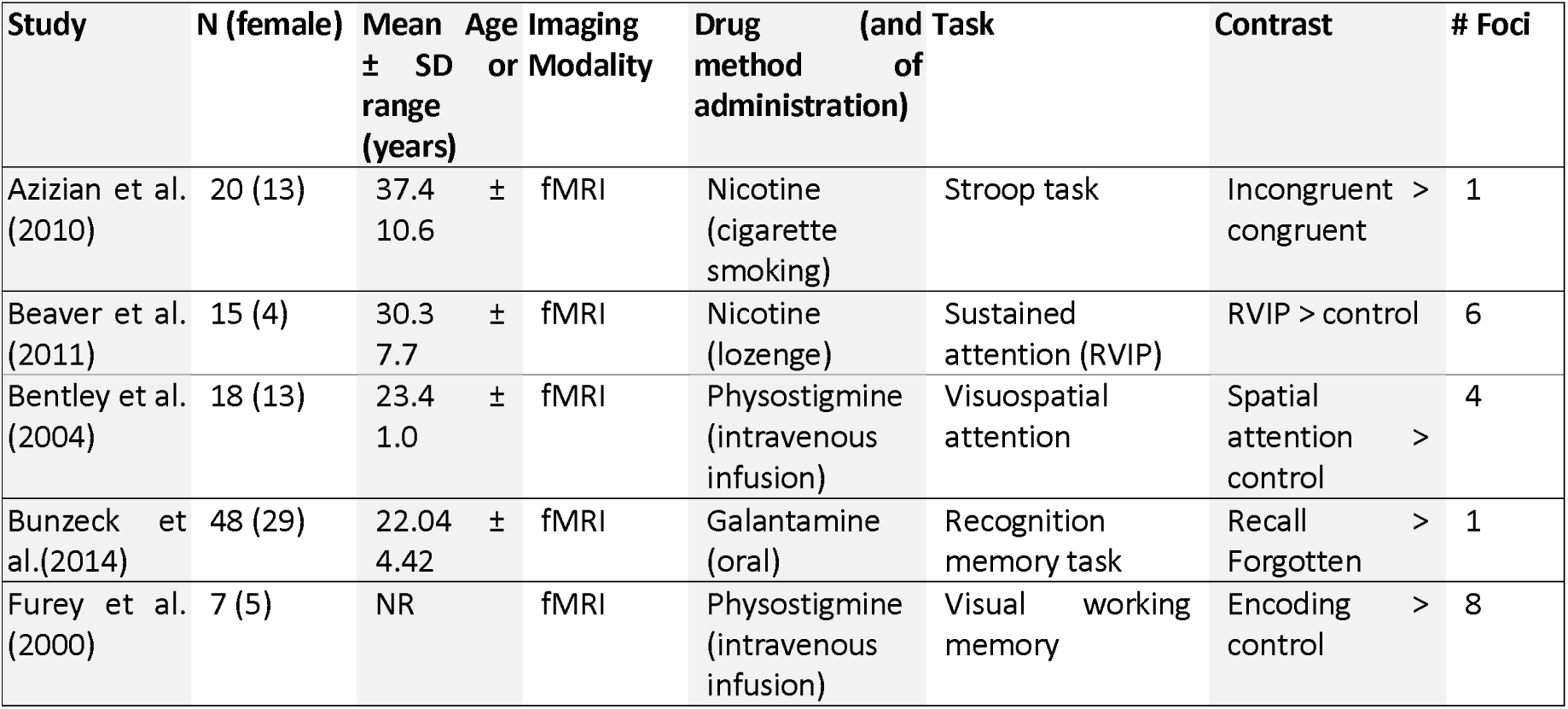
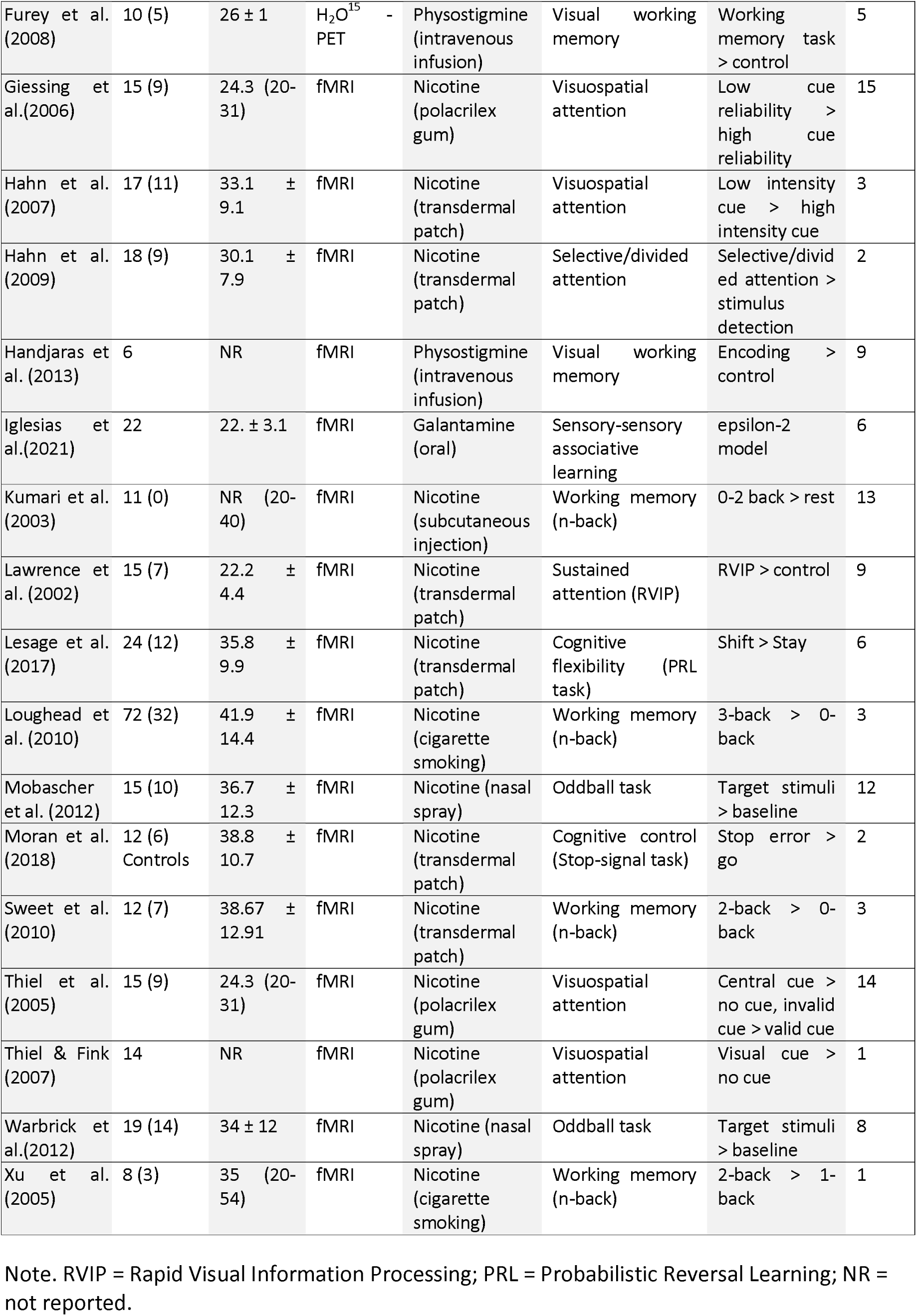
Pharmacological neuroimaging studies reporting increased activation by attention and ACh compared to placebo [ACh-Ag(H>L)]>[Pla(H>L)]

**Table 2.**
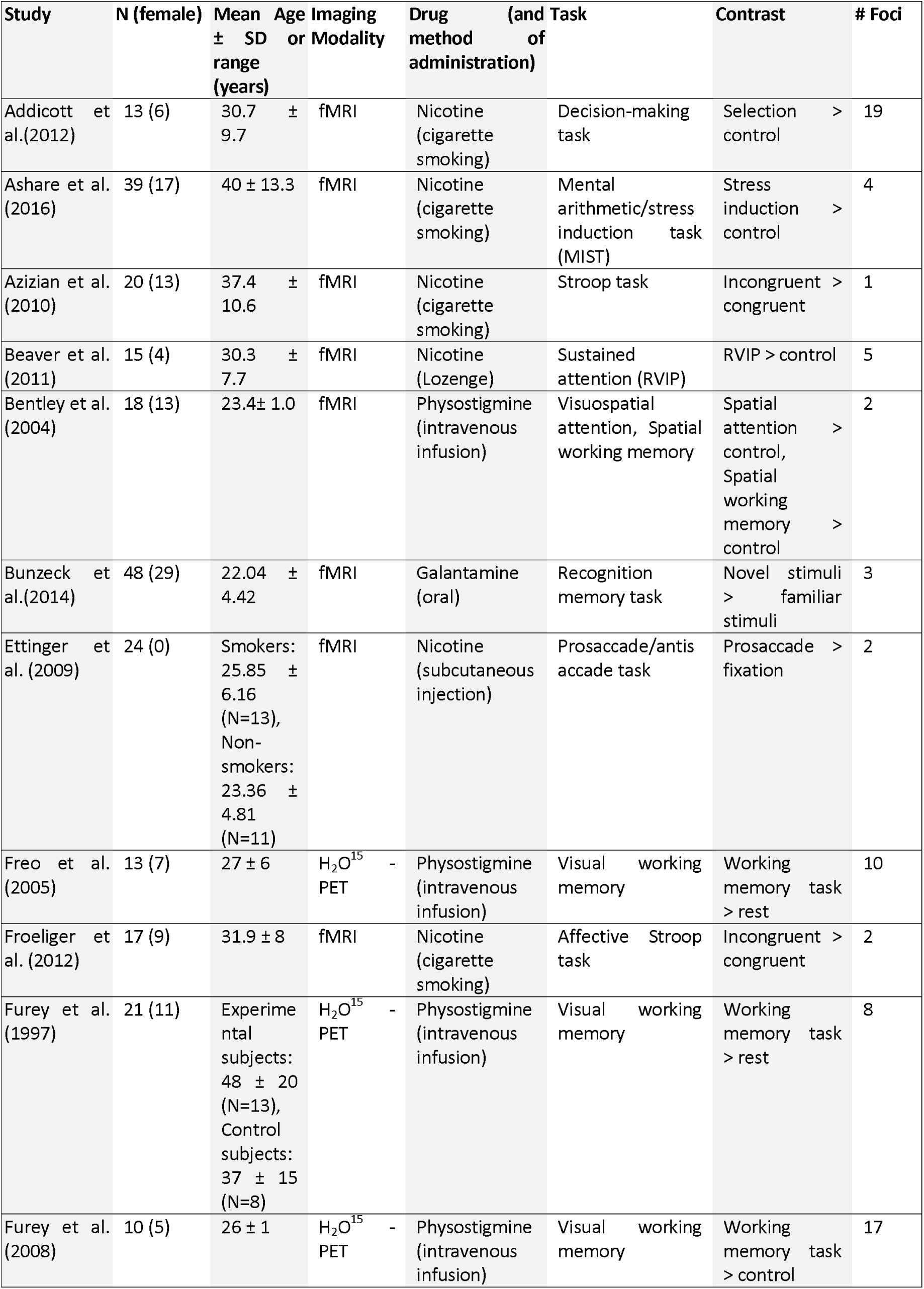
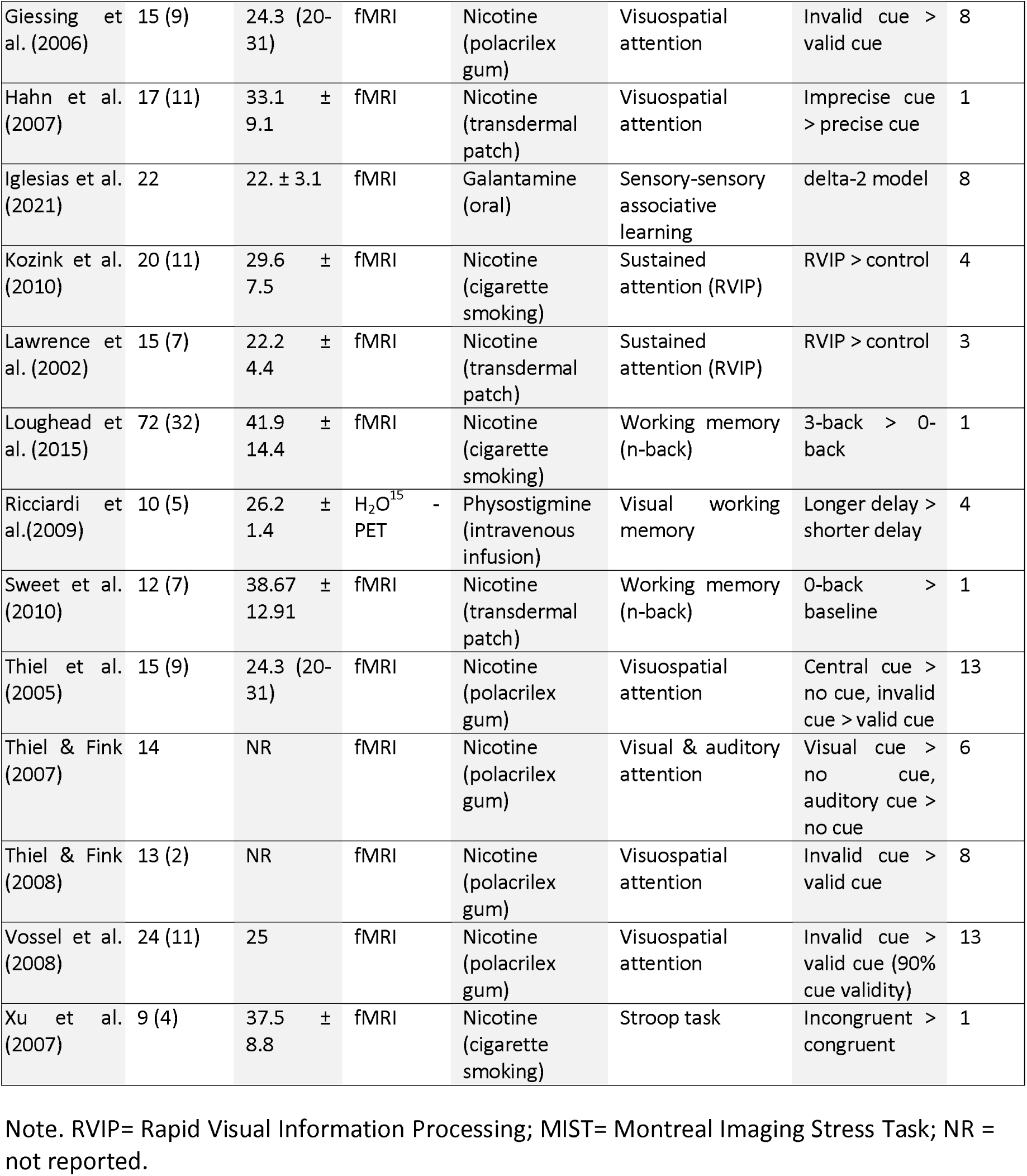
Pharmacological neuroimaging studies reporting decreased activation by attention and ACh compared to placebo [ACh-Ag(H>L)]<[Pla(H>L)].

ALE analyses were performed using GingerALE v.3.0.2 (http://www.brainmap.org/ale/). ALE is a widely used coordinate-based meta-analysis technique which identifies brain regions showing significant convergence in activation across studies. Detailed information regarding ALE and its algorithm has been previously described (Eickhoff *et al*. 2009). Briefly, this approach treats each reported foci as the center of a 3D Gaussian probability distribution rather than as single points in order to account for spatial uncertainty due to between-subject and between-template variance. The width of these distributions is inversely related to the sample size of the corresponding experiment. The probability distribution of all activation foci in each individual contrast is then combined to obtain the modeled activation (MA) maps. ALE maps are subsequently generated by taking the voxel-wise union of these MA maps across all studies. Finally, the computed ALE maps are compared against a null distribution reflecting a random spatial association of experiments to differentiate true convergence of activation foci from random clustering. Following the thresholding guidelines recommended in Eickhoff et al. (2012; 2017), we applied a cluster-level family-wise error (FWE) threshold of *p* > 0.05 with a cluster forming threshold of *p* < 0.001 and 5,000 permutations for each pharmacological imaging ALE analysis.

### Meta-Analytic Connectivity Mapping (MACM)

We performed MACM analyses using Sleuth (https://brainmap.org/sleuth/) separately on two seed regions of interest (ROIs): (1) the *a priori* anatomically defined nuclei of the BF; (2) the suprathreshold clusters identified in the ALE analysis of ACh agonist imaging studies (increases and decreases combined). The binarized MNI space masks for each ROI were entered into the BrainMap database, along with two other search criteria specifying (1) normal mapping and (2) activations. Using the search criteria, MACM queries the database for imaging studies which report coordinates for brain areas that are co-activated with the seed ROI during a particular task or under specific conditions. The coordinates are then entered into an ALE to identify brain regions that are consistently activated across studies.

### Spin tests assessing spatial correspondences among brain maps

Brain maps encoding the ALE Z values were transformed from MNI volume-space to *10k_fsavg* surface-space using neuromaps toolbox (Markello *et al*. 2022) and parcellated using the HCP-MMP 1.0 atlas (Glasser *et al*. 2016). We used a multimodal surface map of the BF connectome derived from (Chakraborty *et al*. 2023); Figure 1A). In neuroimaging studies, spin tests evaluate the statistical significance of spatial relationships between different cortical surface features (Alexander-Bloch *et al*. 2018). Specifically, spin tests can be used to assess whether the observed spatial relationships are significant and unlikely to have occurred by chance. To conduct a spin test on neuroimaging maps, the maps can be randomly permuted or “spun” in a way that maintains the spatial structure of the data but disrupts the relationship between different brain regions. By comparing the observed spatial relationship between two neuroimaging maps to the distribution generated by the spin test, the significance of the relationship can be inferred relative to the spatial null. Spin tests were implemented using the algorithm developed by Alexander-Bloch et al. (2018) to compare cortical maps based on 10,000 permutations.

### Behavioral meta-analysis

We computed separate behavioral meta-analyses for measures of response latency and response accuracy (where available) using the same set of ACh Agonist pharmacological imaging papers reported in the ALE analyses. We used Cohen’s *d* to quantify the standardized mean difference between Drug and Placebo for each behavioral measure. In cases where a measure of effect size other than Cohen’s *d* was reported (e.g. an *F* or *t* statistic), we converted these effect sizes to Cohen’s *d* using formulae specified in Lakens et al (2013) for between group and repeated measures designs. In cases where no measure of effect size was reported, we computed Cohen’s *d* from the means and standard deviations of each group as follows:

For between-group comparisons, Cohen’s *d_s_* was calculated as the difference between the means of the two groups divided by the pooled standard deviation of the two groups. The formula for Cohen’s *d_s_* in this case is:

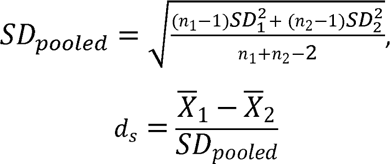

Where *SD_1_* and *SD_2_* are the standard deviations of the two groups being compared, *n_1_* and *n_2_* are the sample sizes of the two groups, *SD_pooled_* is the pooled standard deviation, 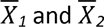 are the means of the two groups being compared.

For repeated measures comparisons, Cohen’s *d_av_* was calculated as the mean difference between the two repeated measures divided by the average standard deviation of both repeated measures. The formula for Cohen’s *d_av_* in this case is:

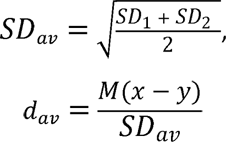

Cohen’s *d_av_* is similar to Cohen’s *d_s_* when correlations between measures are high, and the difference between the standard deviations is low (Lakens 2013). These assumptions are usually met in repeated measures experimental designs where participants perform the same task under two different conditions, e.g., under placebo and ACh.

Cohen’s *d* can provide a biased estimate of the population effect size for studies with smaller sample sizes (*n*<20). Because the majority of studies included in this meta-analysis fall into this smaller size range, we therefore also computed the unbiased Hedge’s *g* effect size estimate from the Cohen’s *d* values above as follows:

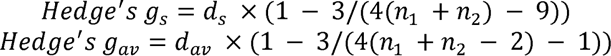

Studies containing *n*>1 statistical comparisons for the effect of ACh on response latency or accuracy, e.g. due to multiple experiments, contributed *n* Cohen’s *d* values to the meta-analyses. Using Cohen’s *d* estimates from each study, we calculated a random-effects weighted average effect size across studies, which took into account both the within- and between-study variance and estimated the precision of this population effect estimate using 95% confidence intervals (Field and Gillett 2010). A random-effects weighted average effect size across studies was also computed on the Hedge’s *g* effect size estimates.

### Data and code availability

All MACM and ALE input data, behavioral meta-analysis input data (Cohen’s *d,* Hedge’s *g*), and code used to conduct the reported spin test analyses and create the figures are available at https://github.com/sudesnac/Ach-phfMRI (Chakraborty).

## Results

### BF co-activates with cortical areas enriched with cholinergic synapses during attention

Extending on our prior work examining the BF cholinergic connectome with multimodal imaging (Chakraborty *et al*. 2023), we first examined which brain areas are consistently co-activated with the BF during attentional engagement. To do so, we employed meta-analytic connectivity mapping (MACM) of the BF (see Methods, Figure 2A). MACM provides a robust and replicable query of brain imaging studies in the BrainMap database (>4,000 studies representing >100,000 participants), returning a list of brain activation coordinates which have been observed to co-occur with activation in *a priori* seed region. Our MACM query returned 88 task imaging experiments reporting BF task-related co-activations in 1357 cognitively normal adults (Figure 2B). We then performed a discovery ALE analysis on the list of brain activation coordinates matching our search criteria to generate a map of the regions that are most likely to be co-activated with BF across experiments. The resulting suprathreshold foci included bilateral anterior insula and frontal opercular cortex, consistent with midcingulo-insular hubs of the ventral attention/salience networks (Figure 3A).

**Figure 3:**
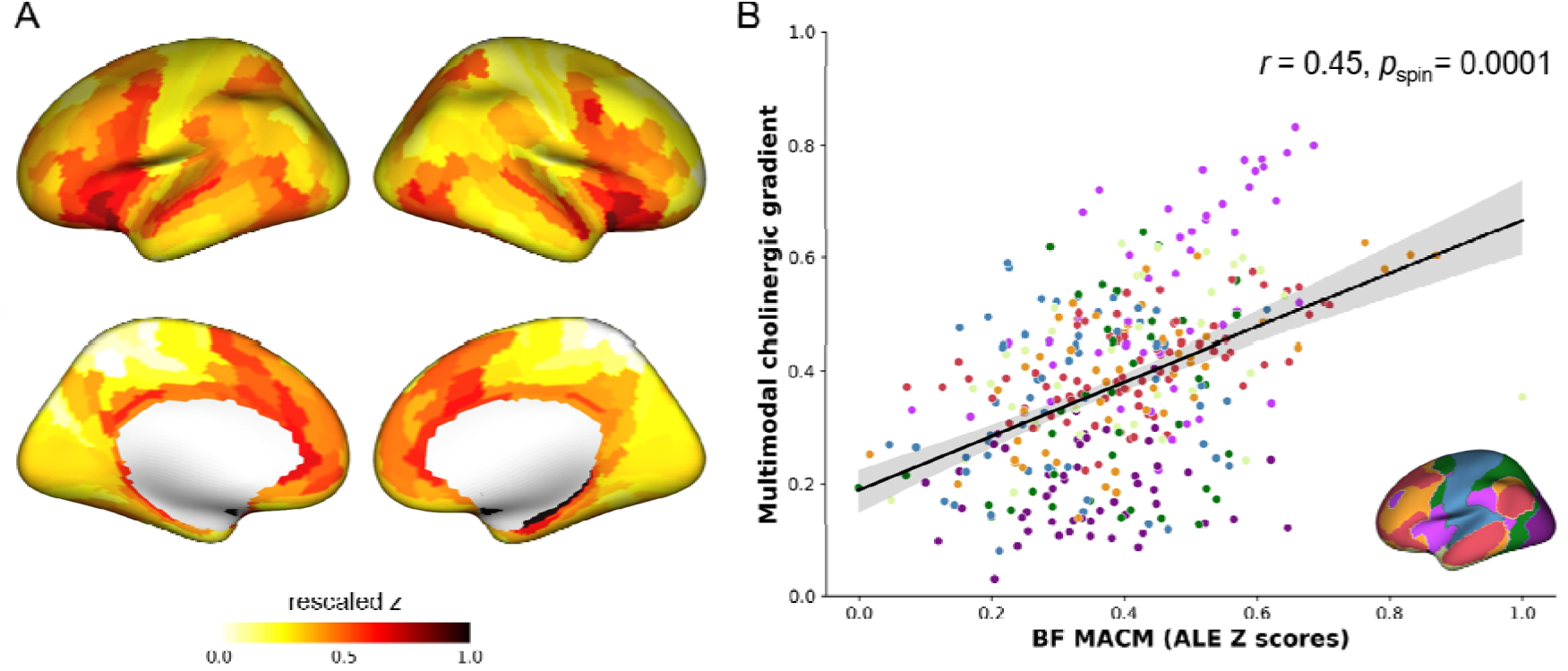
Correlation of BF task co-activation with the multimodal gradient of cortical cholinergic innervation. (A) The ALE Z values for the BF MACM analysis are rescaled 0 to 1 and projected onto the 10k_fsavg cortical surface and parcellated using the HCP-MMP 1.0 atlas. (B) Scatter plot showing the spatial relationship of ALE Z values for the BF MACM (x-axis) with the gradient of cortical cholinergic innervation (y-axis). The r and p-value are derived from a spin test (Alexander-Bloch et al. 2018) between these two surface maps against the spatial null. Each point in the scatter plot represents cortical parcels based on HCP-MMP 1.0 parcellation (Glasser et al. 2016) and is color-coded by the 7 network parcellation from Yeo et al (2011), which is also shown in lower right inset for reference.

We were interested in whether the continuous whole brain map of these ALE effect sizes, encoded by the Z value at each voxel, was spatially related to the multimodal gradient of cortical cholinergic innervation (Chakraborty *et al*. 2023) (Figure 1A). To test this relationship, we transformed the volume map encoding the ALE Z values to the same cortical surface space as the BF cholinergic projectome (Figure 1A). We then performed spin tests against the spatial null to examine their topographical correspondence. The spin test yielded a significant positive correlation (*r*=0.45, *p_spin_*=0.0001), indicating that cortical areas exhibiting the highest task-related co-activations with BF also receive the densest BF cholinergic innervation.

### ACh induces both increased and decreased cingulo-opercular activity during attention

Turning to our core question on the relationship of ACh to brain activity and behavioral performance under attention, we performed meta-analyses on a sample of placebo-controlled fMRI studies examining ACh agonists (e.g. nicotine or acetylcholinesterase inhibitors) in cognitively normal younger adults (<50 years of age) performing experimental tasks containing a manipulation of attentional demand across high (H) and low (L) conditions (see Methods). We identified 33 experiments in total fitting these criteria (Figure 2C). We conducted ALE separately on attention-related activation patterns characterized either by increases under ACh and attention [ACh-Ag(H>L) > Pla(H>L)] (Table 1, *N_studies_*=22, *N_subjects_*=413), or decreases under ACh and attention [ACh-Ag(H>L) < Pla(H>L)] (Table 2, *N_studies_*=24, *N_subjects_*=496). We refer to these patterns as ACh gain and ACh attenuation (Figure 4A).

**Figure 4.**
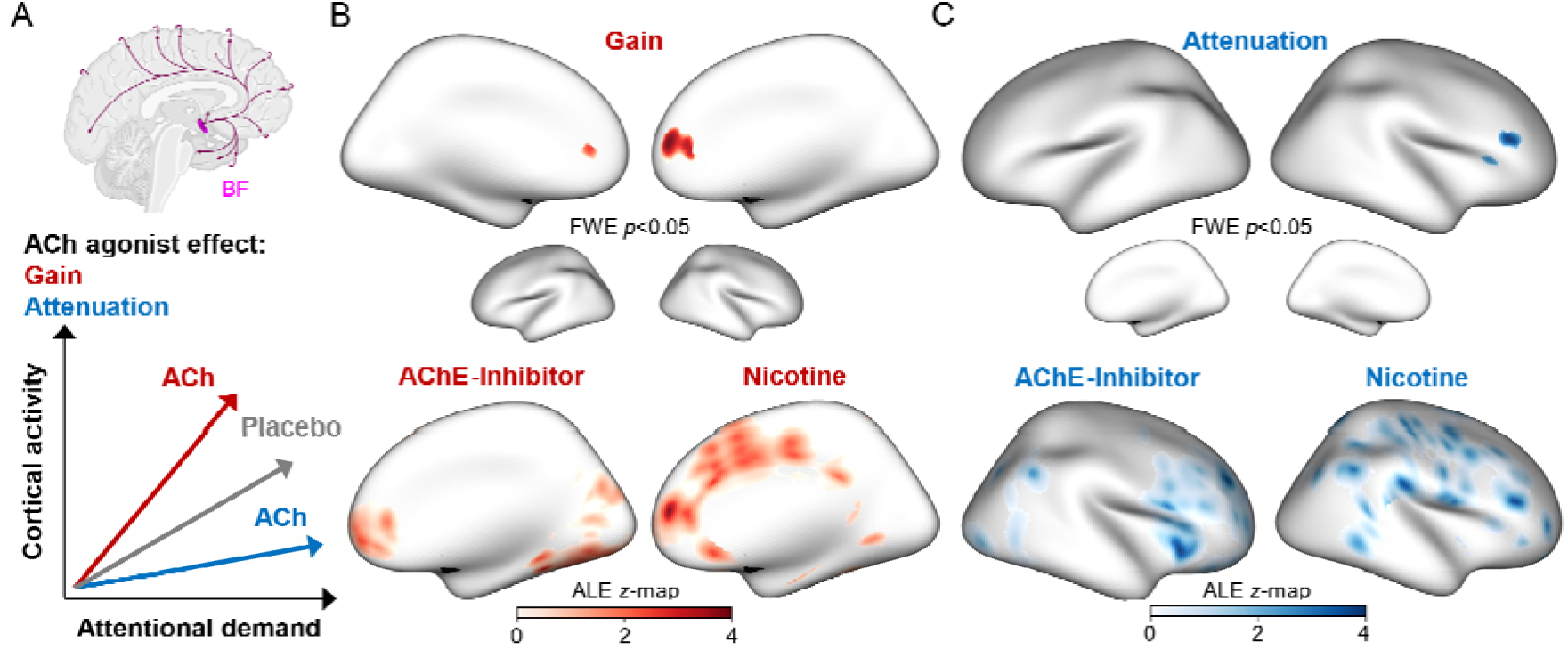
Meta-analyses of task activations under ACh and Attentional demand. (A) Top: Schematic of the basal forebrain (BF) nuclei and its ascending cholinergic projections (magenta). Bottom: Schematic representations of the Drug x Task interactions reported in the literature: Activation gain under ACh agonists is characterized by relatively larger increases in activity with attentional demand compared to placebo (red). Activation attenuation under ACh agonists is characterized by relatively smaller increases in activity with attentional demand compared to placebo. (B) Top row: A significant ACh gain pattern was observed in the anterior cingulate overlapping Brodmann area 13 (cluster level family-wise error FWE corrected p<0.05). Bottom row: Midline surface projections of the ALE z-maps showing the ACh gain effect split according to studies using either nicotine or acetylcholinesterase inhibitors (AChE). (C) Top row: A significant ACh attenuation pattern was observed in the insular and opercular cortex overlapping Brodmann area 32. Bottom row: ALE z-maps for the ACh attenuation effect split according to studies using either nicotine or AChE inhibitors.

The meta-analyses revealed that cholinergic modulation by ACh yielded a distributed pattern of both gain and attenuation of attention-related activity compared to placebo (cluster level FWE corrected *p*<0.05). Areas exhibiting increases under ACh and attention were localized to the right anterior cingulate cortex overlapping Brodmann area 32 (Figure 4B, top rows). By contrast, areas exhibiting decreased activity under attentional demand and ACh were localized to the right opercular and anterior insular cortices overlapping Brodmann area 13 (Figure 4C, top rows).

The above pharmacological studies used one of two categories of cholinergic agonists: nicotine or acetylcholinesterase inhibitors (physostigmine, galantamine). Although these two categories of agonists increase ACh activity, their pharmacological mechanisms of action are not identical (Ishibashi *et al*. 2014; Akaike *et al*. 2010). Our study sample size was underpowered for quantitative comparisons between studies using nicotine and acetylcholinesterase inhibitor agonists (*N_nicotine-increase_*=16, *N_nicotine-decrease_*=17, *N_ACHEI-increase_*=6, *N*_ACHEI-decrease_=7). For comparative visualization, we provide cortical surface projections of the ALE Z values splitting the ACh increases and decreases meta-analyses into nicotine and acetylcholinesterase inhibitor subsamples (Figure 4B and C, bottom panels).

### ACh modulates cortical areas enriched with cholinergic synapses during attention

The meta-analytic observations of activity gain and attenuation under ACh agonists (Figure 4) imply that BF cholinergic projections exert a distributed modulatory effect on right anterior cingulate, opercular and insular cortical areas during directed attention. However, the different activation clusters observed in a standard meta-analysis are not necessarily co-active with one another. For instance, it could be the case that different sets of pharmacological imaging studies in our sample contributed to the significance of each of the observed suprathreshold ALE clusters. We separately confirmed that this was indeed the case; only one experiment reported concurrent activation increases in anterior cingulate and decreases in cingulo-opercular cortex (Furey *et al*. 2008). Are these clusters consistently co-activated with one another under task demand, perhaps reflecting functional embedding within a network, or do they simply reflect coincidental suprathreshold activations loci influenced by different types of experimental tasks? To address this question, we created an ‘ACh’ seed region composed of all suprathreshold clusters exhibiting either gain or attenuation under ACh (Figure 4B,C) and performed a validation MACM and ALE analysis to explicitly test the likelihood of co-activation among these regions across a much larger independent sample of non-pharmacological imaging studies. Using this ACh seed region, along with the same set of additional search criteria used for the BF MACM (constraining to task-related brain activations in normal populations), our query returned 108 experiments, spanning observations from 1530 individuals (Figure 2D). From this sample, we then subjected the activation foci from experiments matching the ACh MACM criteria to ALE analysis of spatial consistency in activity patterns.

We found significant co-activation among the ACh modulated cortical regions during task engagement in this independent sample (Figure 5A). Moreover, extending on the discovery ALE of BF co-activation (Figure 3B), this observed spatial pattern of cortical co-activations provides further validation that these clusters form hubs of the midcingulo-insular network. As with the discovery ALE of BF co-activation, we used spin tests to determine whether the continuous map of ALE Z values encoding the cortical ACh co-activation pattern was also spatially related to the multimodal gradient of cortical cholinergic innervation originating from BF (Chakraborty *et al*. 2023) (Figure 1A). Consistent with a structure-function link between cortical cholinergic innervation and attention-related functional activations, the spin test yielded a significant positive correlation (*r*=0.31, *p_spin_*=0.009), indicating that cortical areas exhibiting the highest task-related co-activations under ACh also receive the densest BF cholinergic innervation (Figure 5B). Altogether, these findings indicate that pharmacological activation of ACh under attentional demand evokes heterogeneous but functionally integrated activation patterns in cortical hubs which overlap the midcingulo-insular hubs of the ventral attention/salience network.

**Figure 5:**
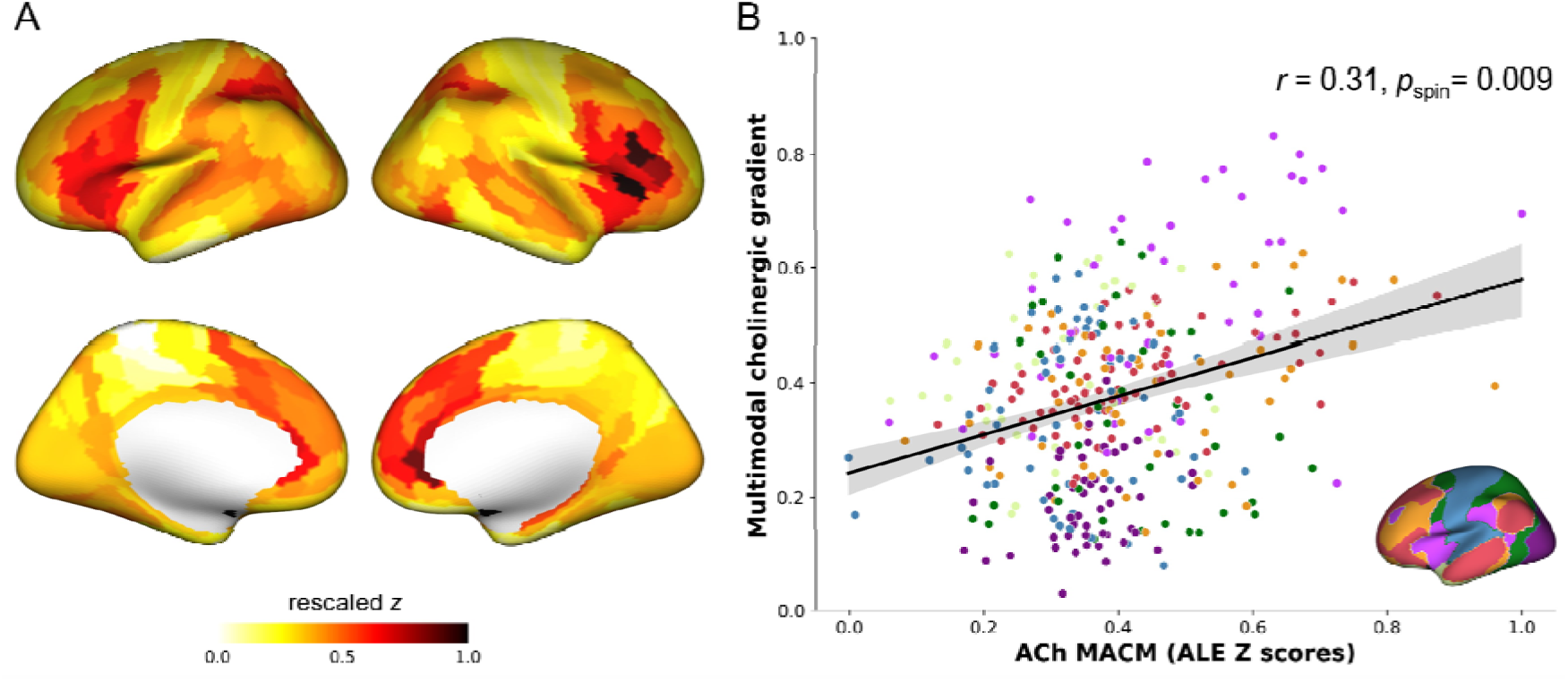
Correlation of ACh cortical co-activation with the multimodal gradient of cortical cholinergic innervation. (A) The ALE Z values for the ACh cortical MACM analysis are rescaled 0 to 1 and projected onto the 10k_fsavg cortical surface and parcellated using the HCP-MMP 1.0 atlas. (B) Scatter plot showing the spatial relationship of ALE Z values for the ACh cortical MACM (x-axis) with the multimodal BF connectome (Figure 1A; y-axis). The r and p-value are derived from a spin test between these two surface maps against a spatial null and is color-coded by the 7 network parcellation from Yeo et al (2011), which is also shown in lower right inset for reference.

### 4.4.4 Modulation of ACh speeds responses with no tradeoff in accuracy

A core inclusion criterion for our meta-analysis of ACh pharmacological fMRI studies was that they employ experimental task manipulations of attentional demand. Of the 33 experiments which met this criterion, some used N-back probes of working memory while others used cue-target selection probes of visuospatial orienting. Although these manipulations probe distinct components of attention, their behavioral correlates are typically reported from common units of response latency and response accuracy, where faster responses in combination with higher or sustained response accuracy, i.e. negligible speed-accuracy tradeoff (Bogacz *et al*. 2010), are thought to reflect superior attentional performance. We therefore computed Cohen’s *d* effect size estimates for comparisons between ACh and placebo (main effect of Drug) on measures of response latency and accuracy. Cohen’s *d* values for each comparison were then submitted to separate random-effects analyses of the weighted average effect size (see Methods, Figure 6).

**Figure 6:**
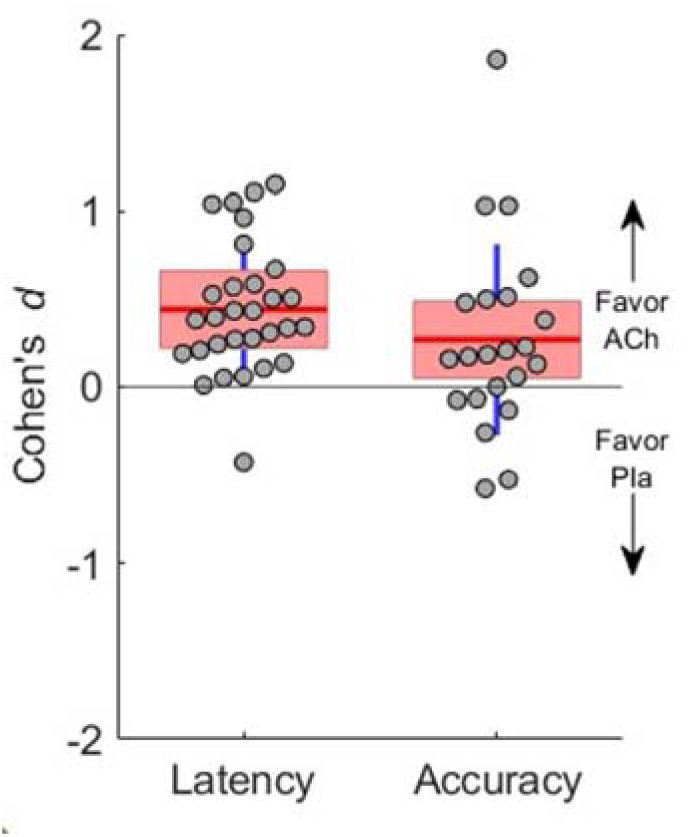
Behavioral meta-analyses for main effects of ACh on response latency and accuracy in pharmacological neuroimaging studies. Cohen’s d values (open circles) estimating the effect of ACh versus placebo for response latencies and response accuracies across experiments. Positive signed Cohen’s d are effect sizes from statistical comparisons which favored performance facilitation under ACh (faster responses, higher accuracy). Negative signed Cohen’s d are effect sizes from statistical comparisons which favored performance facilitation under Placebo. Horizontal lines (red) are the means, boxes (pink) represent one standard deviation from the mean, and vertical lines (blue) are the 95% confidence intervals (CI) derived from a random effects meta-analysis on the weighted Cohen’s d.

For behavioral measures of response latency, we found that ACh yielded significant speeding of target selection compared to placebo (weighted Cohen’s *d*=0.39; *95% CI*=[0.26,0.51]; *z*=5.92; *p*<0.001). In a subset of these studies (*n*=12) which also reported Drug x Task interactions, we also detected a significant speeding of responses by ACh under high compared to low attentional demand (weighted Cohen’s *d*=0.69; *95% CI* =[0.49,0.90]; *z*=6.58; *p*<0.001). For behavioral measures of response accuracy, differences between ACh and placebo were lower but remained significant (weighted Cohen’s *d*=0.24; *95% CI*=[0.01,0.47]; *z*=2.05; *p*=0.04). The number of Drug x Task interactions (*n*=3) reported for response accuracy was insufficient for random effects meta-analysis. Analyses of the unbiased Hedge’s *g* effect size estimate recapitulated the above results (response latency main effect: *g*=0.37, *p*<0.001; response latency interaction: *g*=0.66, *p*<0.001; response accuracy main effect: *g*=0.24, *p=*0.04). Altogether, these behavioral meta-analytic findings imply that ACh facilitates faster decisions about targets without tradeoff in target selection accuracy.

## Discussion

In this study, we explored the relationship between ACh and attention in humans using meta-analytic strategies targeting both activation patterns in pharmacological and non-pharmacological neuroimaging studies, along with pharmacologically induced changes in behavioral response speed and accuracy. Compared to placebo, ACh evoked both gain and attenuation of brain activity primarily in right midcingulo-insular cortical areas. We found that these cortical areas exhibit strong task-related co-activation with one another, and with the BF in large independent meta-analyses of non-pharmacological neuroimaging research. Finally, we show that the cortical topographies of BF co-activation and ACh modulation during directed attention both closely overlap with the multimodal gradient of cortical cholinergic innervation (Chakraborty *et al*. 2023). Concurrent to the influence of ACh on brain activity, we found that individuals exhibit faster target selection, without sacrificing selection accuracy, during directed attention.

There is relatively little intersectionality between lines of imaging research on BF connectivity and ACh pharmacology in humans. As such, there are many gaps in our understanding of how the structural organization of the BF cholinergic projections into the cortex shapes cortico-cortical connectivity, and how cortical ACh release within the BF projectome facilitates directed attention to stimuli, task representations and behavioral responses. To measure intrinsic BF connectivity in the human brain *in vivo*, researchers have typically employed either PET or MRI. For PET studies, the [*^18^*F] FEOBV radiotracer is used to target the vesicular acetylcholine transporter (VAChT), a glycoprotein expressed exclusively by cholinergic neurons (Kanel *et al*. 2022; Albin *et al*. 2018). Because VAChT is expressed most strongly on the presynaptic terminals, these studies have provided insights into the anatomical distribution of the BF cholinergic projections throughout the human cerebrum with cell type specificity. This work is complemented by a growing number of MRI studies examining the white matter and resting state functional connectivity of the BF (Markello *et al*. 2018; Yuan *et al*. 2019; Li *et al*. 2014; Nemy *et al*. 2020; Ray *et al*. 2015; Teipel *et al*. 2011; Lin *et al*. 2022; Oswal *et al*. 2021; Grothe *et al*. 2021; Zhang *et al*. 2017; Fritz *et al*. 2019). We recently employed multimodal PET/MR imaging to examine the spatial relationships among cortical VAChT concentrations, BF white matter projections, and BF resting-state functional connectivity (Chakraborty *et al*. 2023). We demonstrated that cortical areas exhibiting higher VAChT concentrations receive BF projections with greater neuronal branch size and complexity, as estimated by structure-function detethering (Paquola *et al*. 2019; Vázquez-Rodríguez *et al*. 2019; Suárez *et al*. 2020). Moreover, this multimodal map of the BF cortical cholinergic projectome exhibited a striking convergence with midcingulo-insular hubs of the ventral attention and salience networks (Corbetta *et al*. 2008; Vossel *et al*. 2014; Uddin 2015; Uddin *et al*. 2019; Menon and Uddin 2010; Seeley *et al*. 2007; Seeley 2019; Sridharan *et al*. 2008). These findings indicate that cortical ACh signaling plays a key role in allocating attentional resources throughout the brain (see Figure 1). However, pharmacological neuroimaging is needed to determine whether and how ACh modulates cortical function, and how this functional modulation translates to attentional performance. In this study, we therefore used meta-analytic techniques to bridge imaging research on ACh connectivity and ACh pharmacology. We hope that these studies motivate further hypothesis-driven research to strengthen our understanding of the critical roles of ACh in human brain function.

Our meta-analytic findings indicate that with increasing attentional demand, ACh induced both increased activity in the right cingulate cortex and decreased activity in right insular-opercular relative to placebo. This pattern is difficult to interpret from the brain imaging modalities used by studies in our meta-analyses, which included fMRI blood oxygenation level dependent (BOLD) signals and H *^15^*O PET regional cerebral blood flow (rCBF). Neither of these imaging modalities can distinguish excitatory and inhibitory neuronal activity. However, multiple lines of electrophysiological, MRI, histological and optogenetic evidence in non-human animal models indicates that ACh release in the cortex may increase and decrease neuronal activity in parallel across different spatiotemporal scales (Mesulam 2004; Medalla and Barbas 2012; Khalighinejad *et al*. 2020; Galvin *et al*. 2020; Galvin *et al*. 2018; Vijayraghavan *et al*. 2018; Kuchibhotla *et al*. 2017; Do *et al*. 2016; Saunders *et al*. 2015; Chen *et al*. 2012). From this work, ACh appears to exert its modulatory effects on cortical activity primarily via synaptic connections with multiple classes of interneurons and astrocytes, as opposed to direct excitatory synapses with pyramidal neurons (Kuchibhotla *et al*. 2017; Chen *et al*. 2012). These modulatory effects can be inhibitory or disinhibitory, depending on task context and sensory stimuli. The BF is also populated by diverse types of neurons in addition to cholinergic neurons, including long-range projecting glutamatergic neurons and GABAergic interneurons, which regulate cortical states through coordinated activity (Do *et al*. 2016). Adding yet further complexity to these modulatory effects, BF cholinergic neurons are capable of manufacturing and co-releasing ACh and GABA (Saunders *et al*. 2015). The concurrent activity increases and decreases detected by the pharmacological neuroimaging meta-analyses may reflect parallel disinhibitory and inhibitory influences of ACh on midcingulo-insular activity during attention. This study has limitations related to meta-analysis in general and pharmacological neuroimaging in particular. Generally, meta-analysis is susceptible to publication bias and variability in data quality, study populations and experimental designs. We used several study inclusion criteria to mitigate heterogeneity in the populations examined and in the experimental designs employed across studies. A limitation more specific to our meta-analysis of pharmacological neuroimaging studies concerns the relatively small size of our sample, which may increase risk for both Type I and Type II error. To mitigate this limitation, we conducted larger discovery and validation meta-analyses using non-pharmacological imaging studies. Finally, it should be noted that the interrelationships among pharmacological manipulation of ACh, fMRI BOLD, rCBF PET and neuronal activity are indirect. Inferences drawn from these studies about the modulatory effects of ACh on neuronal activity merit caution. The cholinergic system is a complex network of neuronal and receptor subtypes. These include striatal and cortical cholinergic interneurons, in addition to the large projection cholinergic neurons of the BF (Ahmed *et al*. 2019). ACh signaling via nicotinic and muscarinic receptor subtypes is increasingly understood to operate on potentially separate spatiotemporal scales (Obermayer *et al*. 2017). This physiological complexity is difficult to resolve with imaging techniques currently feasible in humans. The cholinergic system is also involved in many critical physiological functions, such as vasodilation, some of which may directly impact the cerebral blood flow component of BOLD and rCBF responses (Hamner *et al*. 2012). Nevertheless, we show that the spatial pattern of brain co-activations identified by our meta-analyses closely resembles areas which are densely innervated by the BF cholinergic projections, assayed by cell type specific PET radiotracers and multimodal MRI measures of BF connectivity.

In sum, the present meta-analytic findings provide further evidence that the midcingulo-insular network is a cortical ‘hotspot’ for BF cholinergic innervation and ACh modulatory activity during attentionally demanding tasks.

## Abbreviations

ACh: acetylcholine
ACh-Ag: acetylcholinergic agonist
AChE: acetylcholinesterase
ALE: activation likelihood estimate
BF: basal forebrain
BOLD: blood oxygen level dependent
FEOBV: fluoroethoxy-benzovesamicol
MA: modelled activation
MACM: meta-analytic connectivity mapping
MRI: magnetic resonance imaging
PET: positron emission tomography
Pla: placebo
rCBF: regional cerebral blood flow
VAChT: vesicular acetylcholine transporter

## Author contributions

SC: Data curation, Formal analysis, Methodology, Resources, Software, Visualization, Writing – original draft, Writing – review & editing. SKL: Data curation, Formal analysis, Methodology, Writing – review & editing. SMA: Data curation, Formal analysis, Methodology, Writing – review & editing. RAMH: Formal analysis, Methodology, Supervision, Visualization, Writing – original draft, Writing – review & editing. ARK: Methodology, Project administration, Resources, Supervision, Writing – original draft, Writing – review & editing. TWS: Conceptualization, Data curation, Formal analysis, Funding acquisition, Investigation, Methodology, Project administration, Resources, Software, Supervision, Validation, Visualization, Writing – original draft, Writing – review & editing.

## Data and code Availability

The configuration files for input to ALE meta-analysis, ALE output maps, spin tests and surface visualizations are available at: https://github.com/sudesnac/Ach-phfMRI. The code for computing ALE maps is available at: https://www.brainmap.org/ale/.

## Reference

Addicott M. A., Baranger D. A. A., Kozink R. V., Smoski M. J., Dichter G. S., McClernon F. J. (2012) Smoking withdrawal is associated with increases in brain activation during decision making and reward anticipation: a preliminary study. Psychopharmacology 219, 563–573.

Ahmed N. Y., Knowles R., Dehorter N. (2019) New Insights Into Cholinergic Neuron Diversity. Front. Mol. Neurosci. 12, 204.

Akaike A., Takada-Takatori Y., Kume T., Izumi Y. (2010) Mechanisms of neuroprotective effects of nicotine and acetylcholinesterase inhibitors: role of alpha4 and alpha7 receptors in neuroprotection. J. Mol. Neurosci. 40, 211–216.

Albin R. L., Bohnen N. I., Muller M. L. T. M., Dauer W. T., Sarter M., Frey K. A., Koeppe R. A. (2018) Regional vesicular acetylcholine transporter distribution in human brain: A [18 F]fluoroethoxybenzovesamicol positron emission tomography study. J. Comp. Neurol. 526, 2884–2897.

Alexander-Bloch A. F., Shou H., Liu S., Satterthwaite T. D., Glahn D. C., Shinohara R. T., Vandekar S. N., Raznahan A. (2018) On testing for spatial correspondence between maps of human brain structure and function. Neuroimage 178, 540–551.

Alves P. N., Forkel S. J., Corbetta M., Thiebaut de Schotten M. (2022) The subcortical and neurochemical organization of the ventral and dorsal attention networks. Commun Biol 5, 1343.

Ashare R. L., Lerman C., Cao W., Falcone M., Bernardo L., Ruparel K., Hopson R., Gur R., Pruessner J. C., Loughead J. (2016) Nicotine withdrawal alters neural responses to psychosocial stress. Psychopharmacology 233, 2459–2467.

Azizian A., Nestor L. J., Payer D., Monterosso J. R., Brody A. L., London E. D. (2010) Smoking reduces conflict-related anterior cingulate activity in abstinent cigarette smokers performing a Stroop task. Neuropsychopharmacology 35, 775–782.

Beaver J. D., Long C. J., Cole D. M., Durcan M. J., Bannon L. C., Mishra R. G., Matthews P. M. (2011) The effects of nicotine replacement on cognitive brain activity during smoking withdrawal studied with simultaneous fMRI/EEG. Neuropsychopharmacology 36, 1792– 1800.

Bennett C., Arroyo S., Berns D., Hestrin S. (2012) Mechanisms generating dual-component nicotinic EPSCs in cortical interneurons. J. Neurosci. 32, 17287–17296.

Bentley P., Driver J., Dolan R. J. (2011) Cholinergic modulation of cognition: insights from human pharmacological functional neuroimaging. Prog. Neurobiol. 94, 360–388.

Bentley P., Husain M., Dolan R. J. (2004) Effects of cholinergic enhancement on visual stimulation, spatial attention, and spatial working memory. Neuron 41, 969–982.

Bogacz R., Wagenmakers E.-J., Forstmann B. U., Nieuwenhuis S. (2010) The neural basis of the speed-accuracy tradeoff. Trends Neurosci. 33, 10–16.

Bunzeck N., Guitart-Masip M., Dolan R. J., Duzel E. (2014) Pharmacological dissociation of novelty responses in the human brain. Cereb. Cortex 24, 1351–1360.

Chakraborty S. Ach-phfMRI: meta-analysis of pharmacological fMRI studies on cholinergic agonists. Github.

Chakraborty S., Haast R. A. M., Kanel P., Khan A. R., Schmitz T. W. (2023) Multimodal gradients of human basal forebrain connectivity.

Chen N., Sugihara H., Sharma J., Perea G., Petravicz J., Le C., Sur M. (2012) Nucleus basalis-enabled stimulus-specific plasticity in the visual cortex is mediated by astrocytes. Proc. Natl. Acad. Sci. U. S. A. 109, E2832–41.

Corbetta M., Patel G., Shulman G. L. (2008) The reorienting system of the human brain: from environment to theory of mind. Neuron 58, 306–324.

Do J. P., Xu M., Lee S.-H., Chang W.-C., Zhang S., Chung S., Yung T. J., et al. (2016) Cell type-specific long-range connections of basal forebrain circuit. Elife 5, e13214.

Downar J., Crawley A. P., Mikulis D. J., Davis K. D. (2000) A multimodal cortical network for the detection of changes in the sensory environment. Nat. Neurosci. 3, 277–283.

Eickhoff S. B., Bzdok D., Laird A. R., Kurth F., Fox P. T. (2012) Activation likelihood estimation meta-analysis revisited. Neuroimage 59, 2349–2361.

Eickhoff S. B., Laird A. R., Fox P. M., Lancaster J. L., Fox P. T. (2017) Implementation errors in the GingerALE Software: Description and recommendations. Hum. Brain Mapp. 38, 7– 11.

Eickhoff S. B., Laird A. R., Grefkes C., Wang L. E., Zilles K., Fox P. T. (2009) Coordinate-based activation likelihood estimation meta-analysis of neuroimaging data: a random-effects approach based on empirical estimates of spatial uncertainty. Hum. Brain Mapp. 30, 2907–2926.

Ettinger U., Williams S. C. R., Patel D., Michel T. M., Nwaigwe A., Caceres A., Mehta M. A., Anilkumar A. P., Kumari V. (2009) Effects of acute nicotine on brain function in healthy smokers and non-smokers: estimation of inter-individual response heterogeneity. Neuroimage 45, 549–561.

Field A. P., Gillett R. (2010) How to do a meta-analysis. Br. J. Math. Stat. Psychol. 63, 665– 694.

Freo U., Ricciardi E., Pietrini P., Schapiro M. B., Rapoport S. I., Furey M. L. (2005) Pharmacological modulation of prefrontal cortical activity during a working memory task in young and older humans: a PET study with physostigmine. Am. J. Psychiatry 162, 2061–2070.

Fritz H.-C. J., Ray N., Dyrba M., Sorg C., Teipel S., Grothe M. J. (2019) The corticotopic organization of the human basal forebrain as revealed by regionally selective functional connectivity profiles. Hum. Brain Mapp. 40, 868–878.

Froeliger B., Modlin L., Wang L., Kozink R. V., McClernon F. J. (2012) Nicotine withdrawal modulates frontal brain function during an affective Stroop task. Psychopharmacology 220, 707–718.

Furey M. L., Pietrini P., Haxby J. V. (2000) Cholinergic enhancement and increased selectivity of perceptual processing during working memory. Science 290, 2315–2319.

Furey M. L., Pietrini P., Haxby J. V., Alexander G. E., Lee H. C., VanMeter J., Grady C. L., et al. (1997) Cholinergic stimulation alters performance and task-specific regional cerebral blood flow during working memory. Proc. Natl. Acad. Sci. U. S. A. 94, 6512–6516.

Furey M. L., Ricciardi E., Schapiro M. B., Rapoport S. I., Pietrini P. (2008) Cholinergic enhancement eliminates modulation of neural activity by task difficulty in the prefrontal cortex during working memory. J. Cogn. Neurosci. 20, 1342–1353.

Galvin V. C., Arnsten A. F. T., Wang M. (2018) Evolution in Neuromodulation—The Differential Roles of Acetylcholine in Higher Order Association vs. Primary Visual Cortices. Front. Neural Circuits 12.

Galvin V. C., Yang S. T., Paspalas C. D., Yang Y., Jin L. E., Datta D., Morozov Y. M., et al. (2020) Muscarinic M1 Receptors Modulate Working Memory Performance and Activity via KCNQ Potassium Channels in the Primate Prefrontal Cortex. Neuron 106, 649–661.e4.

Giessing C., Thiel C. M., Rösler F., Fink G. R. (2006) The modulatory effects of nicotine on parietal cortex activity in a cued target detection task depend on cue reliability. Neuroscience 137, 853–864.

Glasser M. F., Coalson T. S., Robinson E. C., Hacker C. D., Harwell J., Yacoub E. (2016) A multi-modal parcellation of human cerebral cortex. Nature Publishing Group 536, 171–178.

Grothe M. J., Labrador-Espinosa M. A., Jesús S., Macías-García D., Adarmes-Gómez A., Carrillo F., Camacho E. I., et al. (2021) In vivo cholinergic basal forebrain degeneration and cognition in Parkinson’s disease: Imaging results from the COPPADIS study. Parkinsonism Relat. Disord. 88, 68–75.

Guo W., Robert B., Polley D. B. (2019) The Cholinergic Basal Forebrain Links Auditory Stimuli with Delayed Reinforcement to Support Learning. Neuron 103, 1164–1177.e6.

Hahn B., Ross T. J., Wolkenberg F. A., Shakleya D. M., Huestis M. A., Stein E. A. (2009) Performance effects of nicotine during selective attention, divided attention, and simple stimulus detection: an fMRI study. Cereb. Cortex 19, 1990–2000.

Hahn B., Ross T. J., Yang Y., Kim I., Huestis M. A., Stein E. A. (2007) Nicotine enhances visuospatial attention by deactivating areas of the resting brain default network. J. Neurosci. 27, 3477–3489.

Hamner J. W., Tan C. O., Tzeng Y.-C., Taylor J. A. (2012) Cholinergic control of the cerebral vasculature in humans. J. Physiol. 590, 6343–6352.

Handjaras G., Ricciardi E., Szczepanik J., Pietrini P., Furey M. L. (2013) Cholinergic enhancement differentially modulates neural response to encoding during face identity and face location working memory tasks. Exp. Biol. Med. 238, 999–1008.

Hangya B., Ranade S. P., Lorenc M., Kepecs A. (2015) Central Cholinergic Neurons Are Rapidly Recruited by Reinforcement Feedback. Cell 162, 1155–1168.

Harrison T. C., Pinto L., Brock J. R., Dan Y. (2016) Calcium Imaging of Basal Forebrain Activity during Innate and Learned Behaviors. Front. Neural Circuits 10, 36.

Hasselmo M. E., McGaughy J. (2004) High acetylcholine levels set circuit dynamics for attention and encoding and low acetylcholine levels set dynamics for consolidation. Prog. Brain Res. 145, 207–231.

Iglesias S., Kasper L., Harrison S. J., Manka R., Mathys C., Stephan K. E. (2021) Cholinergic and dopaminergic effects on prediction error and uncertainty responses during sensory associative learning. Neuroimage 226, 117590.

Ishibashi M., Yamazaki Y., Miledi R., Sumikawa K. (2014) Nicotinic and muscarinic agonists and acetylcholinesterase inhibitors stimulate a common pathway to enhance GluN2B-NMDAR responses. Proc. Natl. Acad. Sci. U. S. A. 111, 12538–12543.

Jing M., Li Y., Zeng J., Huang P., Skirzewski M., Kljakic O., Peng W., et al. (2020) An optimized acetylcholine sensor for monitoring in vivo cholinergic activity. Nat. Methods.

Kanel P., Zee S. van der, Sanchez-Catasus C. A., Koeppe R. A., Scott P. J. H., Laar T. van, Albin R. L., Bohnen N. I. (2022) Cerebral topography of vesicular cholinergic transporter changes in neurologically intact adults: A [18F]FEOBV PET study. Aging Brain 2, 100039.

Khalighinejad N., Bongioanni A., Verhagen L., Folloni D., Attali D., Aubry J.-F., Sallet J., Rushworth M. F. S. (2020) A Basal Forebrain-Cingulate Circuit in Macaques Decides It Is Time to Act. Neuron 105, 370–384.e8.

Kozink R. V., Lutz A. M., Rose J. E., Froeliger B., McClernon F. J. (2010) Smoking withdrawal shifts the spatiotemporal dynamics of neurocognition. Addict. Biol. 15, 480–490.

Kuchibhotla K. V., Gill J. V., Lindsay G. W., Papadoyannis E. S., Field R. E., Sten T. A. H., Miller K. D., Froemke R. C. (2017) Parallel processing by cortical inhibition enables context-dependent behavior. Nat. Neurosci. 20, 62–71.

Kumari V., Gray J. A., Ffytche D. H., Mitterschiffthaler M. T., Das M., Zachariah E., Vythelingum G. N., Williams S. C. R., Simmons A., Sharma T. (2003) Cognitive effects of nicotine in humans: an fMRI study. Neuroimage 19, 1002–1013.

Lakens D. (2013) Calculating and reporting effect sizes to facilitate cumulative science: a practical primer for t-tests and ANOVAs. Front. Psychol. 4, 863.

Lancaster J. L., Tordesillas-Gutiérrez D., Martinez M., Salinas F., Evans A., Zilles K., Mazziotta J. C., Fox P. T. (2007) Bias between MNI and Talairach coordinates analyzed using the ICBM-152 brain template. Hum. Brain Mapp. 28, 1194–1205.

Laszlovszky T., Schlingloff D., Hegedüs P., Freund T. F., Gulyás A., Kepecs A., Hangya B. (2020) Distinct synchronization, cortical coupling and behavioral function of two basal forebrain cholinergic neuron types. Nat. Neurosci. 23, 992–1003.

Lawrence N. S., Ross T. J., Stein E. A. (2002) Cognitive mechanisms of nicotine on visual attention. Neuron 36, 539–548.

Lesage E., Aronson S. E., Sutherland M. T., Ross T. J., Salmeron B. J., Stein E. A. (2017) Neural Signatures of Cognitive Flexibility and Reward Sensitivity Following Nicotinic Receptor Stimulation in Dependent Smokers: A Randomized Trial. JAMA Psychiatry 74, 632–640.

Letzkus J. J., Wolff S. B. E., Meyer E. M. M., Tovote P., Courtin J., Herry C., Lüthi A. (2011) A disinhibitory microcircuit for associative fear learning in the auditory cortex. Nature 480, 331–335.

Li C.-S. R., Ide J. S., Zhang S., Hu S., Chao H. H., Zaborszky L. (2014) Resting state functional connectivity of the basal nucleus of Meynert in humans: in comparison to the ventral striatum and the effects of age. Neuroimage 97, 321–332.

Lin C. P., Frigerio I., Boon B. D. C., Zhou Z., Rozemuller A. J. M., Bouwman F. H., Schoonheim M. M., Berg W. D. J. van de, Jonkman L. E. (2022) Structural (dys)connectivity associates with cholinergic cell density in Alzheimer’s disease. Brain 145, 2869–2881.

Li X., Yu B., Sun Q., Zhang Y., Ren M., Zhang X., Li A., et al. (2018) Generation of a whole-brain atlas for the cholinergic system and mesoscopic projectome analysis of basal forebrain cholinergic neurons. Proc. Natl. Acad. Sci. U. S. A. 115, 415–420.

Loughead J., Ray R., Wileyto E. P., Ruparel K., Sanborn P., Siegel S., Gur R. C., Lerman C. (2010) Effects of the α4β2 Partial Agonist Varenicline on Brain Activity and Working Memory in Abstinent Smokers. Biol. Psychiatry 67, 715–721.

Loughead J., Wileyto E. P., Ruparel K., Falcone M., Hopson R., Gur R., Lerman C. (2015) Working memory-related neural activity predicts future smoking relapse. Neuropsychopharmacology 40, 1311–1320.

Mandino F., Vrooman R. M., Foo H. E., Yeow L. Y., Bolton T. A. W., Salvan P., Teoh C. L., et al. (2022) A triple-network organization for the mouse brain. Mol. Psychiatry 27, 865–872.

Markello R. D., Hansen J. Y., Liu Z.-Q., Bazinet V., Shafiei G., Suárez L. E., Blostein N., et al. (2022) neuromaps: structural and functional interpretation of brain maps. Nat. Methods.

Markello R. D., Spreng R. N., Luh W.-M., Anderson A. K., De Rosa E. (2018) Segregation of the human basal forebrain using resting state functional MRI. Neuroimage 173, 287– 297.

Medalla M., Barbas H. (2012) The anterior cingulate cortex may enhance inhibition of lateral prefrontal cortex via m2 cholinergic receptors at dual synaptic sites. J. Neurosci. 32, 15611–15625.

Menon V., Uddin L. Q. (2010) Saliency, switching, attention and control: a network model of insula function. Brain Struct. Funct. 214, 655–667.

Mesulam M. M. (2004) The cholinergic innervation of the human cerebral cortex. Prog. Brain Res. 145, 67–78.

Mesulam M. -M, Geula C. (1988) Nucleus basalis (Ch4) and cortical cholinergic innervation in the human brain: Observations based on the distribution of acetylcholinesterase and choline acetyltransferase. J. Comp. Neurol. 275, 216–240.

Mobascher A., Warbrick T., Brinkmeyer J., Musso F., Stoecker T., Jon Shah N., Winterer G. (2012) Nicotine effects on anterior cingulate cortex in schizophrenia and healthy smokers as revealed by EEG-informed fMRI. Psychiatry Res. 204, 168–177.

Moran L. V., Stoeckel L. E., Wang K., Caine C. E., Villafuerte R., Calderon V., Baker J. T., et al. (2018) Nicotine-induced activation of caudate and anterior cingulate cortex in response to errors in schizophrenia. Psychopharmacology 235, 789–802.

Nemy M., Cedres N., Grothe M. J., Muehlboeck J. S., Lindberg O., Nedelska Z., Stepankova O., et al. (2020) Cholinergic white matter pathways make a stronger contribution to attention and memory in normal aging than cerebrovascular health and nucleus basalis of Meynert. Neuroimage 211, 116607.

Obermayer J., Verhoog M. B., Luchicchi A., Mansvelder H. D. (2017) Cholinergic Modulation of Cortical Microcircuits Is Layer-Specific: Evidence from Rodent, Monkey and Human Brain. Front. Neural Circuits 11, 100.

Oswal A., Gratwicke J., Akram H., Jahanshahi M., Zaborszky L., Brown P., Hariz M., Zrinzo L., Foltynie T., Litvak V. (2021) Cortical connectivity of the nucleus basalis of Meynert in Parkinson’s disease and Lewy body dementias. Brain 144, 781–788.

Paquola C., Vos De Wael R., Wagstyl K., Bethlehem R. A. I., Hong S.-J., Seidlitz J., Bullmore E. T., et al. (2019) Microstructural and functional gradients are increasingly dissociated in transmodal cortices. PLoS Biol. 17, e3000284.

Pinto L., Goard M. J., Estandian D., Xu M., Kwan A. C., Lee S. H., Harrison T. C., Feng G., Dan Y. (2013) Fast modulation of visual perception by basal forebrain cholinergic neurons. Nat. Neurosci. 16, 1857–1863.

Ray N. J., Metzler-Baddeley C., Khondoker M. R., Grothe M. J., Teipel S., Wright P., Heinsen H., Jones D. K., Aggleton J. P., O’Sullivan M. J. (2015) Cholinergic basal forebrain structure influences the reconfiguration of white matter connections to support residual memory in mild cognitive impairment. J. Neurosci. 35, 739–747.

Ricciardi E., Pietrini P., Schapiro M. B., Rapoport S. I., Furey M. L. (2009) Cholinergic modulation of visual working memory during aging: a parametric PET study. Brain Res. Bull. 79, 322–332.

Sarter M., Lustig C. (2020) Forebrain Cholinergic Signaling: Wired and Phasic, Not Tonic, and Causing Behavior. J. Neurosci. 40, 712–719.

Sarter M., Parikh V., Howe W. M. (2009) Phasic acetylcholine release and the volume transmission hypothesis: time to move on. Nat. Rev. Neurosci. 10, 383–390.

Saunders A., Granger A. J., Sabatini B. L. (2015) Corelease of acetylcholine and GABA from cholinergic forebrain neurons. Elife 4, e06412.

Schmitz T. W., Duncan J. (2018) Normalization and the Cholinergic Microcircuit: A Unified Basis for Attention. Trends Cogn. Sci. 22, 422–437.

Seeley W. W. (2019) The Salience Network: A Neural System for Perceiving and Responding to Homeostatic Demands. J. Neurosci. 39, 9878–9882.

Seeley W. W., Menon V., Schatzberg A. F., Keller J., Glover G. H., Kenna H., Reiss A. L., Greicius M. D. (2007) Dissociable intrinsic connectivity networks for salience processing and executive control. J. Neurosci. 27, 2349–2356.

Sethuramanujam S., Matsumoto A., deRosenroll G., Murphy-Baum B., Grosman C., McIntosh J. M., Jing M., et al. (2021) Rapid multi-directed cholinergic transmission in the central nervous system. Nat. Commun. 12, 1374.

Sforazzini F., Schwarz A. J., Galbusera A., Bifone A., Gozzi A. (2014) Distributed BOLD and CBV-weighted resting-state networks in the mouse brain. Neuroimage 87, 403–415.

Sridharan D., Levitin D. J., Menon V. (2008) A critical role for the right fronto-insular cortex in switching between central-executive and default-mode networks. Proc. Natl. Acad. Sci. U. S. A. 105, 12569–12574.

Suárez L. E., Markello R. D., Betzel R. F., Misic B. (2020) Linking Structure and Function in Macroscale Brain Networks. Trends Cogn. Sci. 24, 302–315.

Sutherland M. T., Ray K. L., Riedel M. C., Yanes J. A., Stein E. A., Laird A. R. (2015) Neurobiological impact of nicotinic acetylcholine receptor agonists: an activation likelihood estimation meta-analysis of pharmacologic neuroimaging studies. Biol. Psychiatry 78, 711–720.

Sweet L. H., Mulligan R. C., Finnerty C. E., Jerskey B. A., David S. P., Cohen R. A., Niaura R. S. (2010) Effects of nicotine withdrawal on verbal working memory and associated brain response. Psychiatry Res. 183, 69–74.

Teipel S. J., Meindl T., Grinberg L., Grothe M., Cantero J. L., Reiser M. F., Moller H. J., Heinsen H., Hampel H. (2011) The cholinergic system in mild cognitive impairment and Alzheimer’s disease: an in vivo MRI and DTI study. Hum. Brain Mapp. 32, 1349–1362.

Teles-Grilo Ruivo L. M., Baker K. L., Conway M. W., Kinsley P. J., Gilmour G., Phillips K. G., Isaac J. T. R., Lowry J. P., Mellor J. R. (2017) Coordinated Acetylcholine Release in Prefrontal Cortex and Hippocampus Is Associated with Arousal and Reward on Distinct Timescales. Cell Rep. 18, 905–917.

Thiel C. M., Fink G. R. (2007) Visual and auditory alertness: modality-specific and supramodal neural mechanisms and their modulation by nicotine. J. Neurophysiol. 97, 2758–2768.

Thiel C. M., Fink G. R. (2008) Effects of the cholinergic agonist nicotine on reorienting of visual spatial attention and top-down attentional control. Neuroscience 152, 381–390.

Thiel C. M., Zilles K., Fink G. R. (2005) Nicotine modulates reorienting of visuospatial attention and neural activity in human parietal cortex. Neuropsychopharmacology 30, 810–820.

Tu G., Halawa A., Yu X., Gillman S., Takehara-Nishiuchi K. (2022) Outcome-Locked Cholinergic Signaling Suppresses Prefrontal Encoding of Stimulus Associations. J. Neurosci. 42, 4202–4214.

Uddin L. Q. (2015) Salience processing and insular cortical function and dysfunction. Nat. Rev. Neurosci. 16, 55–61.

Uddin L. Q., Yeo B. T. T., Spreng R. N. (2019) Towards a Universal Taxonomy of Macro-scale Functional Human Brain Networks. Brain Topogr. 32, 926–942.

Vázquez-Rodríguez B., Suárez L. E., Markello R. D., Shafiei G., Paquola C., Hagmann P., Heuvel M. P. van den, Bernhardt B. C., Spreng R. N., Misic B. (2019) Gradients of structure–function tethering across neocortex. Proceedings of the National Academy of Sciences 116, 21219–21227.

Vijayraghavan S., Major A. J., Everling S. (2018) Muscarinic M1 Receptor Overstimulation Disrupts Working Memory Activity for Rules in Primate Prefrontal Cortex. Neuron 98, 1256–1268.e4.

Vossel S., Geng J. J., Fink G. R. (2014) Dorsal and ventral attention systems: distinct neural circuits but collaborative roles. Neuroscientist 20, 150–159.

Vossel S., Thiel C. M., Fink G. R. (2008) Behavioral and neural effects of nicotine on visuospatial attentional reorienting in non-smoking subjects. Neuropsychopharmacology 33, 731–738.

Warbrick T., Mobascher A., Brinkmeyer J., Musso F., Stoecker T., Shah N. J., Fink G. R., Winterer G. (2012) Nicotine effects on brain function during a visual oddball task: a comparison between conventional and EEG-informed fMRI analysis. J. Cogn. Neurosci. 24, 1682–1694.

Woolf N. J. (1991) Cholinergic systems in mammalian brain and spinal cord. Prog. Neurobiol. 37, 475–524.

Wu H., Williams J., Nathans J. (2014) Complete morphologies of basal forebrain cholinergic neurons in the mouse. Elife 3, e02444.

Xu J., Mendrek A., Cohen M. S., Monterosso J., Rodriguez P., Simon S. L., Brody A., et al. (2005) Brain activity in cigarette smokers performing a working memory task: effect of smoking abstinence. Biol. Psychiatry 58, 143–150.

Xu J., Mendrek A., Cohen M. S., Monterosso J., Simon S., Jarvik M., Olmstead R., Brody A. L., Ernst M., London E. D. (2007) Effect of cigarette smoking on prefrontal cortical function in nondeprived smokers performing the Stroop Task. Neuropsychopharmacology 32, 1421–1428.

Xu N., LaGrow T. J., Anumba N., Lee A., Zhang X., Yousefi B., Bassil Y., et al. (2022) Functional Connectivity of the Brain Across Rodents and Humans. Front. Neurosci. 16, 816331.

Yeo B. T. T., Krienen F. M., Sepulcre J., Sabuncu M. R., Lashkari D., Hollinshead M., Roffman J. L., et al. (2011) The organization of the human cerebral cortex estimated by intrinsic functional connectivity. J. Neurophysiol. 106, 1125–1165.

Yeung A. W. K., Robertson M., Uecker A., Fox P. T., Eickhoff S. B. (2023) Trends in the sample size, statistics, and contributions to the BrainMap database of activation likelihood estimation meta-analyses: An empirical study of 10-year data. Hum. Brain Mapp. 44, 1876–1887.

Yuan R., Biswal B. B., Zaborszky L. (2019) Functional Subdivisions of Magnocellular Cell Groups in Human Basal Forebrain: Test–Retest Resting-State Study at Ultra-high Field, and Meta-analysis. Cereb. Cortex 29, 2844–2858.

Zaborszky L., Duque A., Gielow M., Gombkoto P., Nadasdy Z., Somogyi J. (2015) Organization of the Basal Forebrain Cholinergic Projection System: Specific or Diffuse? The Rat Nervous System: Fourth Edition, 491–507.

Záborszky L., Gombkoto P., Varsanyi P., Gielow M. R., Poe G., Role L. W., Ananth M., et al. (2018) Specific Basal Forebrain–Cortical Cholinergic Circuits Coordinate Cognitive Operations. J. Neurosci. 38, 9446–9458.

Zaborszky L., Hoemke L., Mohlberg H., Schleicher A., Amunts K., Zilles K. (2008) Stereotaxic probabilistic maps of the magnocellular cell groups in human basal forebrain. Neuroimage 42, 1127–1141.

Zhang S., Hu S., Fucito L. M., Luo X., Mazure C. M., Zaborszky L., Li C.-S. R. (2017) Resting-State Functional Connectivity of the Basal Nucleus of Meynert in Cigarette Smokers: Dependence Level and Gender Differences. Nicotine Tob. Res. 19, 452–459.

